# Estimating Gait Kinematics from Muscle Activity Using Deep Learning in Typically Developing Children

**DOI:** 10.64898/2026.02.05.703957

**Authors:** Carmen Fernández-González, Beatriz De la Calle, Carlos Gómez, Hichem Saoudi, Daniel Iordanov, Francesco Cenni, Mario Martínez-Zarzuela

**Author notes:** Corresponding autor.

## Abstract

Instrumented gait assessment in pediatric populations is often constrained by the complexity and lack of portability of traditional motion capture systems. In this article, we propose a deep learning approach utilizing a one-dimensional (1D) U-Net architecture to accurately estimate ankle and knee joint angles in the sagittal plane from surface electromyography (sEMG) signals. We analyzed data from the tibialis anterior and medial gastrocnemius of 25 typically developing children (ages 4–16) to evaluate the model’s performance and the influence of age-related gait maturation. The proposed 1D U-Net achieved high predictive accuracy for the ankle joint (RMSE: 3.6°) and the knee joint (RMSE: 4.1∘). Experimental results demonstrated that incorporating the toe-off event as a temporal marker significantly enhanced prediction stability during transitional gait phases. Furthermore, Statistical Parametric Mapping (SPM) was employed to identify systematic errors, which were primarily localized during initial contact and pre-swing but remained below clinically relevant thresholds. The findings reveal that prediction accuracy increases with age, reflecting more stable neuromotor patterns. This study demonstrates that a 1D U-Net can reliably decode lower-limb kinematics from sEMG alone, enabling the development of simplified, non-invasive, and portable pediatric gait assessment tools that can be integrated into the control strategies of assistive devices.

## 1. Introduction

Gait analysis is a key tool for understanding human locomotion, offering insights into motor control, musculoskeletal dynamics, and functional performance (Akhtaruzzaman et al., 2016). It plays a central role in clinical assessment and rehabilitation, as well as in biomechanics research. Despite advances in motion analysis technology (Bonato et al., 2024), obtaining accurate gait kinematics often requires complex and costly laboratory setups, which limits its accessibility in routine practice (Rodrigues et al., 2019).

Traditional gait evaluation combines observational and instrumental methods. Visual scales, such as the *Edinburgh Visual Gait Score*, provide a qualitative evaluation of kinematic deviations, while clinical instruments, like the *Modified Ashworth Scale*, offer a subjective assessment of muscular spasticity (McGibbon et al., 2013). On the other hand, instrumental methods, such as optoelectronic motion capture systems (*e*.*g*., *Vicon*), allow a more objective and detailed characterization of movement. Despite their accuracy, motion capture systems require complex and costly infrastructures, including dedicated laboratory spaces, multiple infrared cameras, and reflective markers, which limit their accessibility in clinical contexts (Rodrigues et al., 2019). To overcome these constraints, inertial measurement units (IMUs) and surface electromyography (sEMG) have emerged as more portable alternatives (Cimolin & Galli, 2014),(Huang et al., 2022). Nevertheless, IMU-based assessments still depend on multi-sensor setups and calibration procedures that can be time-consuming. While sensor placement is generally less intrusive than in the past, obtaining high-quality signals requires well-trained personnel, and data quality may still be compromised by suboptimal sensor positioning or calibration. These limitations have driven the need for methods capable of inferring kinematic information from more accessible sources without the need for additional sensors.

Within this context, understanding gait in the pediatric population represents a particularly relevant challenge. Unlike adults, children exhibit continuous changes in gait patterns as a result of musculoskeletal growth, motor learning, and neuromuscular maturation (Dewolf et al., 2020). Gait is not considered to reach an adult-like form until approximately eight years of age, when spatiotemporal and kinematic parameters begin to stabilize (Froehle et al., 2013; Sutherland, 1997). Establishing normative references for typical developing children is therefore crucial for developmental studies and also for the early detection of motor impairments.

Recent research has demonstrated the feasibility of predicting joint kinematics from sEMG signals using advanced deep learning models. For instance, one study reported root-mean-square error (RMSE) values of 5.6° at the knee and 2.6° at the ankle, when combining sEMG from lower-limb and trunk muscles (Guez et al., 2025). Another study achieved RMSE values below 8° for the knee using a deep learning–based coupled predictor (Wei et al., 2021). Additionally, models based on Random Forests have obtained errors as low as 3.9° at the ankle (Hollinger et al., 2023),(Amrani El Yaakoubi et al., 2023). Despite these promising results, recent systematic reviews highlight that machine learning applications in this field still face significant hurdles for clinical implementation (Pan et al., 2026). Most of these studies share similar limitations: (i) only adult populations were analyzed, without valid extrapolation to pediatric cohorts; (ii) the experimental conditions were restricted to artificial setups, such as treadmill walking; and (iii) they rely on a large number of sEMG sensors, which limits their clinical applicability.

To address these gaps, the present study offers three main contributions: (i) the development of a methodology capable of estimating ankle and knee joint angles using only two sensors per joint; (ii) the design of an automatic synchronization and segmentation framework for sEMG and IMU signals; and (iii) the identification of a developmental trend showing that, in the absence of gait pathologies, model prediction error decreases with age, providing novel insights into motor maturation processes during childhood. Together, these contributions advance the development of accessible and clinically applicable tools for pediatric gait assessment.

## 2. Materials and Methods

### 2.1. Participants

Twenty-five pediatric participants without gait abnormalities (7.6 ± 2.3 years; 56% male; 68% right laterality) were included in this study. Participants were stratified into < 8 years (*n* = 16) and ≥ 8 years (*n* = 9), based on developmental gait stages (Froehle et al., 2013; Sutherland, 1997). The study was approved by the Research Ethics Committee with Medicines (CEIM) West Valladolid Health Area, in March 2023 (reference protocol 22-PI125), and written informed consent was obtained from all participants and their legal guardians.

### 2.2. Data Collection

Recordings were conducted in the Rehabilitation Unit of Hospital Universitario Río Hortega (Valladolid, Spain) under standardized clinical conditions. Prior to data acquisition, all IMUs were calibrated using the proprietary *Xsens* software. Participants performed the gait analysis in a 12-meter room, starting from a marked point. Each trial began with the participant standing still for 5 seconds, followed by walking to the end of the room and back at a self-selected pace. This sequence was repeated twice per subject to ensure data consistency.

sEMG and kinematic signals were recorded simultaneously. Four *Trigno Avanti* sensors (*Delsys*) captured sEMG activity, while eight *Xsens Awinda* IMUs collected kinematics. Both systems enable the acquisition of angular velocity and acceleration, offering complementary dynamic information. Sensor placement followed standardized anatomical guidelines to ensure reproducibility. sEMG electrodes were positioned on the tibialis anterior (TA) and medial gastrocnemius (MG) of both legs according to SENIAM recommendations (Hermens et al., 2000). IMUs were placed at the L3–L5 spinal region, mid-lateral thigh, upper lateral surface of the lower leg, sternum, and dorsum of each foot, following *Xsens* guidelines.

### 2.3. Database Preprocessing

Basic preprocessing was applied to minimize noise and enhance the extraction of relevant features from the recorded signals.

For sEMG signals, the following steps were performed (Chowdhury et al., 2013): (i) band-pass filtering: a fourth-order Butterworth filter (20–400 Hz) was applied to remove motion artifacts and high-frequency noise; (ii) rectification: the filtered signal was converted to absolute values, ensuring all amplitudes were positive; and (iii) envelope extraction: a smoothed signal was obtained by applying a second-order low-pass Butterworth filter with a cutoff frequency of 6 Hz, thus generating the RMS envelope. For angular velocity signals, an offset correction was applied by subtracting the mean from each channel. No additional filtering was applied to the kinematic data, in line with previous studies indicating that, under controlled acquisition conditions, raw IMU signals are sufficiently reliable for gait analysis without further preprocessing (Mobbs et al., 2022).

### 2.4. Synchronization and segmentation of sEMG and kinematic signals

Due to the lack of access to commercial synchronization software, a custom procedure was implemented to temporally align the sEMG and kinematic data, ensuring that each step identified in the kinematic signal precisely matched its corresponding step in the sEMG signal. The synchronization procedure was based on the angular velocity signals recorded on each lower leg by the sEMG sensor placed over the tibialis anterior and the IMU positioned on the anterolateral aspect of the tibia, assuming that anatomical proximity between both would result in similar angular dynamics. It was performed using the angular velocity around the longitudinal axis of the tibia, corresponding to the X-axis of the *Delsys* IMU and the Y-axis of the *Xsens* unit. Although the sensor coordinate systems differ, both axes represent the same anatomical direction when the devices are aligned on the tibial segment. This axis was selected because it captures the predominant rotational component of the lower leg during movement.

Since the *Delsys* and *Xsens* systems operated at different sampling rates (148.14 Hz and 100 Hz, respectively), the *Delsys* data were downsampled to 100 Hz to enable point-by-point comparison with the *Xsens* signals. Downsampling was performed by adjusting the number of samples in each signal proportionally to the new sampling rate, preserving the temporal structure and key features of the original movement.

The turning point was automatically detected using a moving variance filter applied to the angular velocity signal. Different window sizes were tested, starting from 100 samples and progressively decreasing to identify the optimal value. Considering the *Xsens* sampling rate and the average movement cycle duration (∼1 s), a 20-sample window (equivalent to 200 ms) provided sufficient temporal resolution to detect local variations in angular velocity while maintaining stability against noise. The selected parameters were consistent with previous studies (Farfán et al., 2010; Abbaspour et al., 2020).

Once the turn was detected in both signals, all local maxima in the angular velocity curves were identified. To synchronize them, the maximum value before and after the turn were used as reference points. Ultimately, the position of the reference points was optimized by minimizing the RMSE.

Following synchronization, the signals were segmented to identify heel strike (HS) and toe off (TO) events. This process combines the detection of local maxima and minima in the angular velocity signal. Specifically, all local maxima were first identified, and from each of them, the adjacent minima were located: the minimum preceding the peak was interpreted as the TO event, and the following minimum as the HS. To validate the alignment, the mean error, standard deviation, and correlation coefficient between the signals were calculated.

As the study involved pediatric population, it was necessary to account for the fact that many participants had not yet developed a mature gait pattern. To address this, a step filtering procedure was applied to retain only gait cycles that were consistent and representative. First, all detected steps within each trial were indexed. The first two steps and last two ones were excluded, as these are commonly influenced by initial acceleration and final deceleration. Steps surrounding the turning point (specifically the two preceding and two following) were also removed due to their instability. Finally, a homogeneity filter was applied: the average of the remaining steps was calculated, and any step from the sEMG or IMU signal that deviated by more than 0.30 times the standard deviation from this mean was discarded.

On average, approximately 13 steps per subject and per repetition (two trials) were retained for analysis. This yielded a total of 650 steps used for model training and validation.

### 2.5. Predictive model developing

To estimate joint angles from multichannel sEMG signals, a one dimensional U-Net architecture was designed, consisting of an encoder, a bottleneck, and a decoder. The encoder extracts hierarchical features and compresses the information, while the decoder reconstructs the temporal resolution to generate continuous joint angle predictions. Skip connections between encoder and decoder blocks help preserve low-level structural information. The network receives as input individual gait steps segmented into 101 samples, corresponding to the normalized time course of a single step (0–100%) of sEMG signals from TA and MG. For each input step, the network outputs the corresponding joint angle (knee or ankle), also normalized to 101 samples across the step duration. Figure 1 provides a detailed diagram of this architecture, including the tensor dimensions and operations performed at each stage of the network.

**Figure 1:**
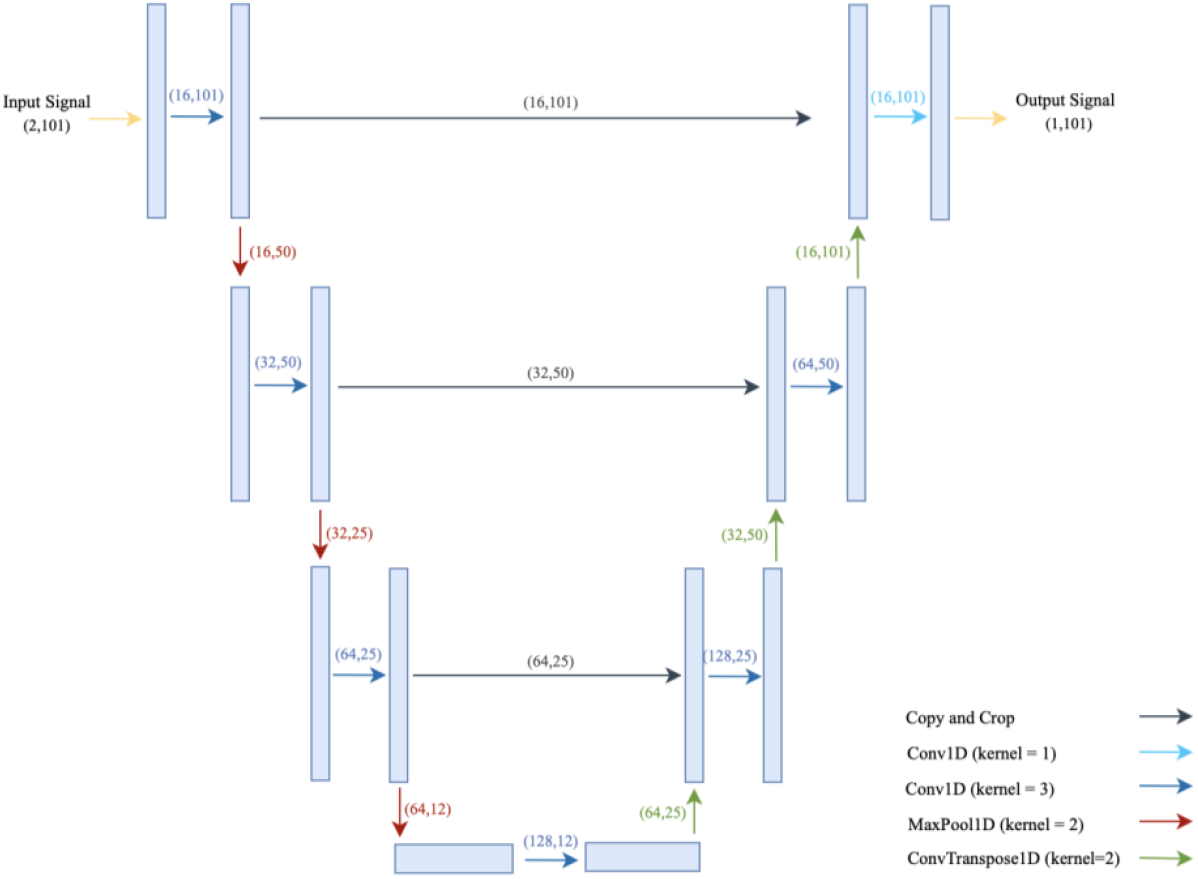
Schematic Representation of the 1D U-Net Architecture Used for Estimating Joint Angles from Multichannel sEMG Input. The diagram illustrates the encoder-decoder structure with convolutional, pooling, and upsampling layers, as well as skip connections for feature preservation.

An experimental variant of the model incorporated an additional input channel: a binary signal indicating the timing of TO event within each window, detected according to the procedure described above. This temporal context was introduced because muscle activations with similar shapes can have different functional roles depending on the phase of the gait cycle (Lacquaniti et al., 2012). By providing explicit information about the TO event, the model can better interpret these activations and improve prediction accuracy, particularly during transitional phases of gait.

The dataset was divided at the subject level into training (70%), validation (15%), and testing (15%) subsets to ensure independence between sets. The model was trained using the *Adam* optimizer, with *dropout* and *early stopping* to prevent overfitting, and an adaptive learning rate schedule to improve convergence. To evaluate different modeling approaches, several experimental configurations were tested, including ankle and knee predictions and models with and without the TO signal. Additionally, predictions were analyzed according to limb dominance to investigate potential differences between dominant and non-dominant limbs. Model performance was assessed using RMSE, MAE, and R^2^.

### 2.6. Temporal Analysis using Statistical Parametric Mapping

In addition to standard performance metrics, it was critical to assess whether model errors occur systematically at specific phases of the gait cycle. To this end, Statistical Parametric Mapping (SPM) was employed, enabling point-by-point comparisons between predicted and reference joint angle trajectories and the identification of temporal regions with significant discrepancies (Pataky et al., 2013).

For each experimental condition, a one-dimensional paired t-test (1D t-test) was applied. Mean predicted and reference trajectories were computed by averaging across repetitions and participants, and a t-statistic was calculated at each of the 101 time points. Cycle segments in which differences exceeded the predefined significance threshold (α = 0.05) were interpreted as systematic deviations in model predictions.

To assess the magnitude and practical relevance of the discrepancies between predicted and reference joint angles, effect sizes were computed using Cohen’s *d*, defined as the mean difference divided by the standard deviation (Diener, 2010). This metric quantifies the strength of the deviation independently of sample size, allowing distinction between statistically detectable and practically meaningful differences. Values of *d* were interpreted according to conventional thresholds (small < 0.5, medium = 0.5–0.8, large > 0.8). In parallel, a clinical relevance threshold was established at 1% of the joint range of motion (ROM) observed during the gait cycle, following previous biomechanical studies (Berner et al., 2020), (Haji Hassani et al., 2022). This corresponds approximately to 2° for the ankle (ROM ≈ 25°) and 5° for the knee (ROM ≈ 65°). Deviations smaller than this limit (typically associated with small effect sizes d < 0.5) were interpreted as biomechanically negligible, even when statistically significant.

## 3. Results

### 3.1. Synchronization and segmentation

Temporal errors for gait events were 0.043 ± 0.021s for HS and 0.049 ± 0.024s for TO. Correlation coefficients between synchronized signals were consistently close to 1.0. Given the 100 Hz sampling rate, these errors represent less than 1% of a typical gait cycle.Representative examples of synchronized sEMG and IMU signals, along with the extracted gait steps, are presented in Figure 2.

**Figure 1:**
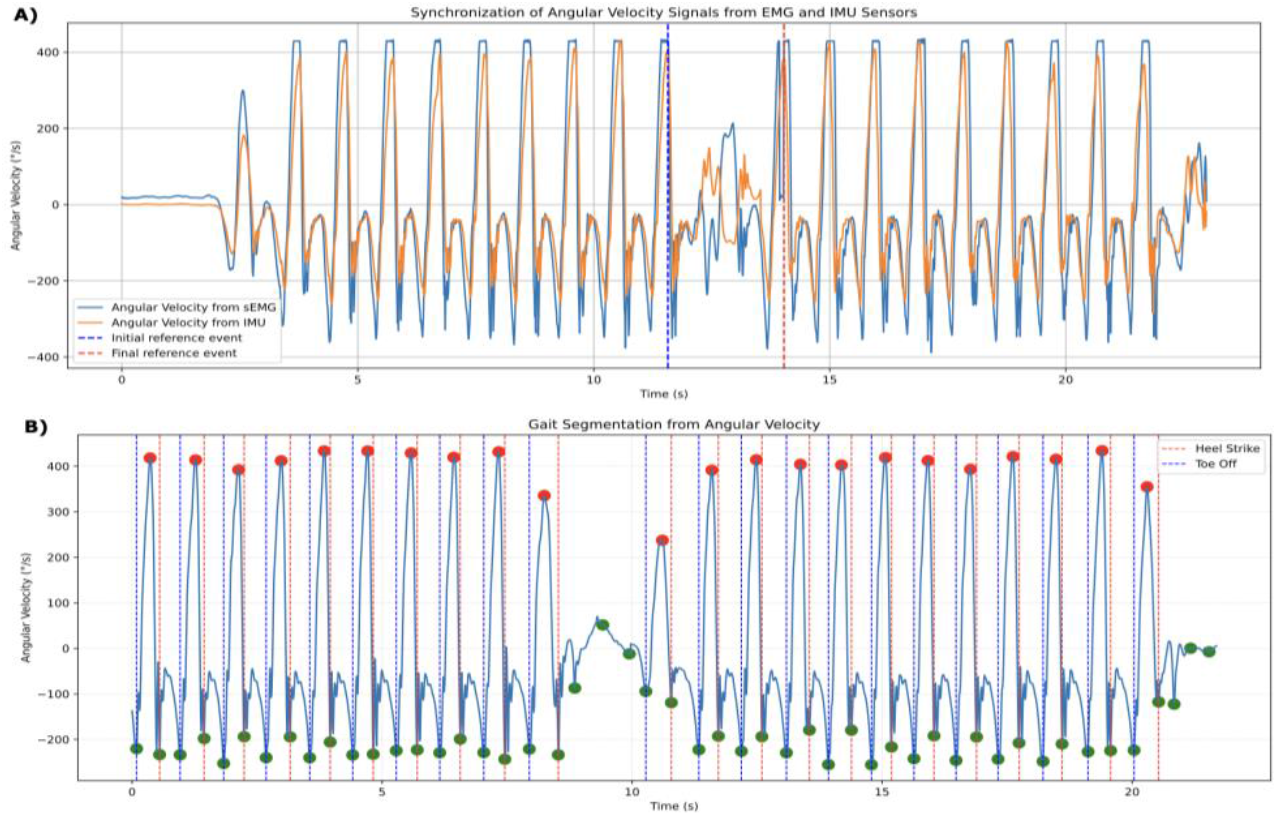
**A)** Synchronization of angular velocity signals from sEMG and IMU sensors; **B)** Gait cycle segmentation based on heel strike and toe-off events.

### 3.2. Prediction for ankle and knee

Table 1 presents the performance of the model across the different configurations, including predictions for the ankle and knee joints, with or without the inclusion of the TO signal, and separated by dominant and non-dominant limbs.

**Table 1:** Overall results of the 1D U-Net model for ankle and knee estimation. RMSE, MAE, and R^2^ are shown for different configurations (TO = Toe Off.

| Configuration | RMSE (°) | MAE (°) | R<sup>2</sup> |
| --- | --- | --- | --- |
| Ankle | 4.9 | 5.1 | 0.8 |
| Ankle + TO | 3.2 | 3.8 | 0.9 |
| Ankle: dominant side + TO | 3.6 | 2.9 | 0.8 |
| Ankle: no dominant side + TO | 3.9 | 3.8 | 0.7 |
| Knee | 6.0 | 5.8 | 0.6 |
| Knee + TO | 4.1 | 3.9 | 0.8 |
| Knee: dominant side + TO | 4.6 | 3.6 | 0.7 |
| Knee: no dominant side + TO | 5.0 | 4.1 | 0.7 |

Figure 3 displays the predicted and reference joint trajectories for all test subjects at both the ankle and the knee. Across all participants, the predicted curves align closely with the reference signals, reproducing the characteristic pattern and amplitude of each gait cycle with remarkable consistency.

**Figure 3.**
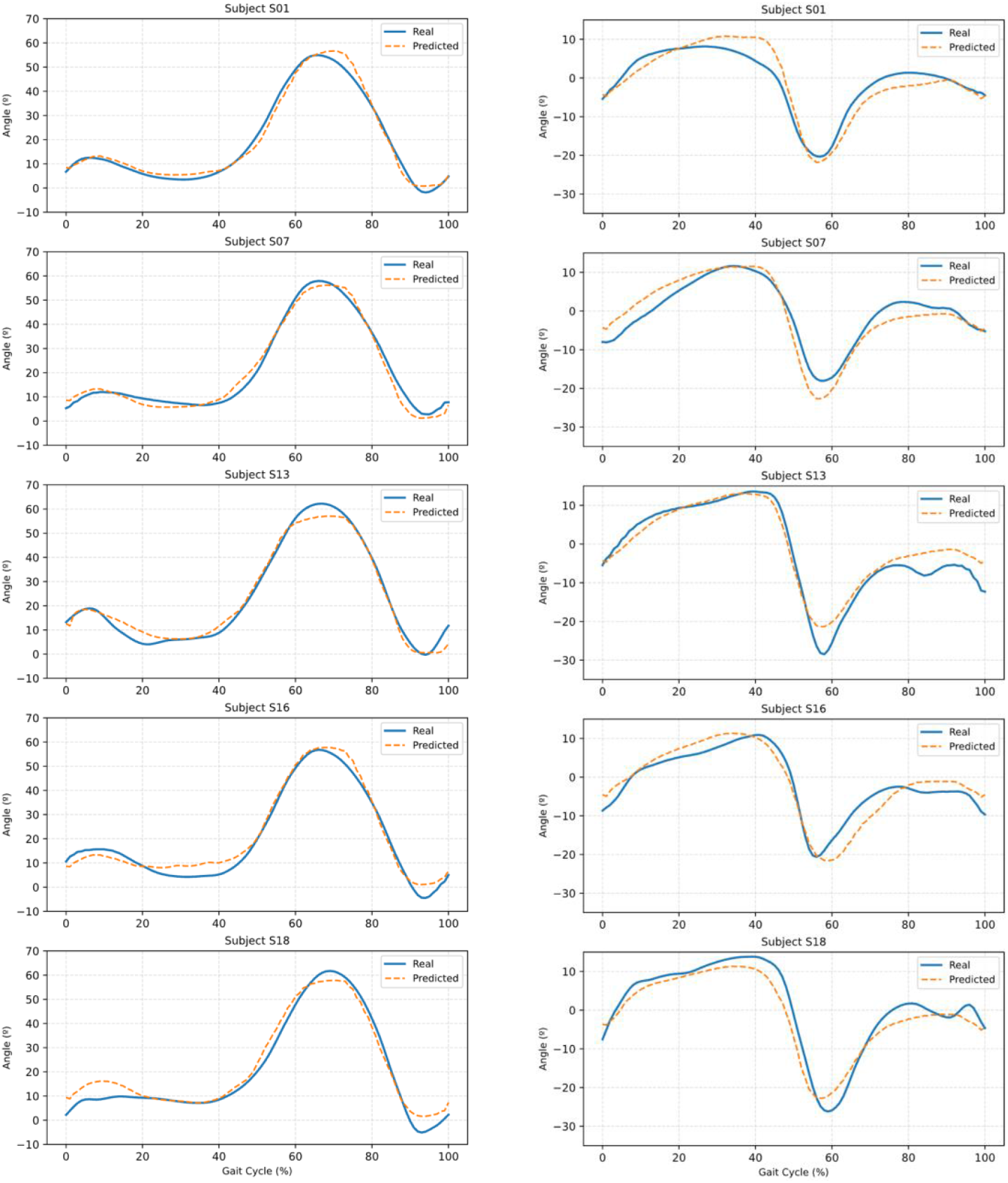
Comparison between predicted and reference joint angle trajectories for the ankle and knee in all subjects from the test group.

### 3.3. SPM analysis

The mean predicted and reference trajectories for the ankle and knee joints are shown in Figure 4, which also includes the results of the SPM analysis. Shaded regions indicate intervals where statistically significant differences (p < 0.05) were detected. A summary of these significant intervals is also provided, showing the temporal location of the deviations for ankle and knee models, with and without TO events.

**Figure 4:**
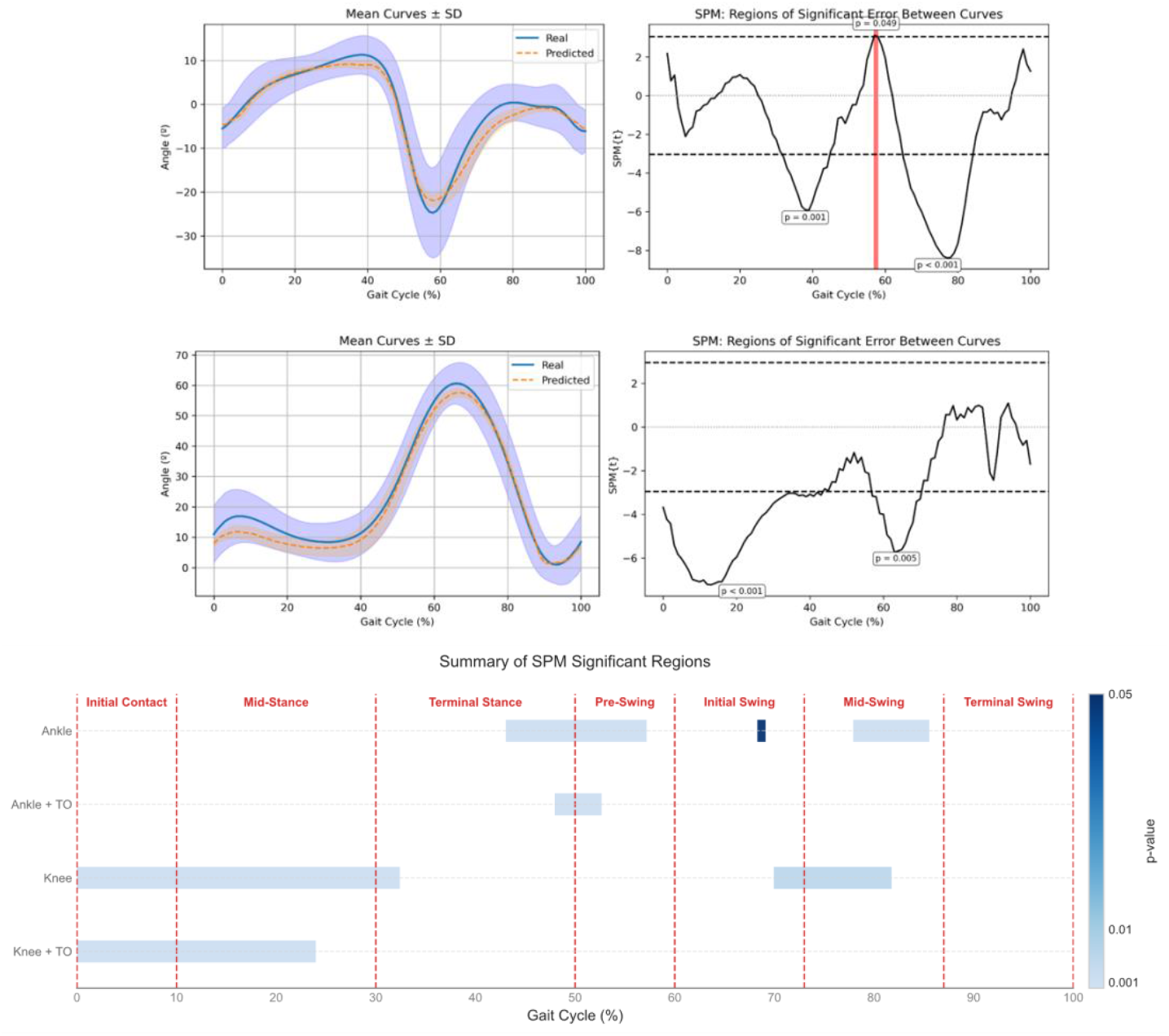
**A)** Mean joint angle trajectories and SPM analysis. Average ± SD of predicted and reference joint angles across the gait cycle. Shaded regions indicate phases where Statistical Parametric Mapping (SPM) detected significant differences (p < 0.05). **B)** Summary of significant intervals for the ankle and knee models, with and without inclusion of toe-off events.

The global effect sizes (Cohen’s d) for each joint, calculated across the full gait cycle, were 0.21 for the ankle and 0.29 for the knee, both classified as small. This indicates that, although statistically detectable differences occur at specific phases of the gait cycle, the overall magnitude of the deviations is minor relative to the variability of the data.

### 3.4. Influence of age

Age-related effects on prediction accuracy were also examined. Figure 5 presents the training and test RMSE values across ages, showing that training errors were consistently and considerably lower than test errors. Overall, both sets of errors decreased with increasing age, reflecting the maturation of gait patterns around eight years.

**Figure 5:**
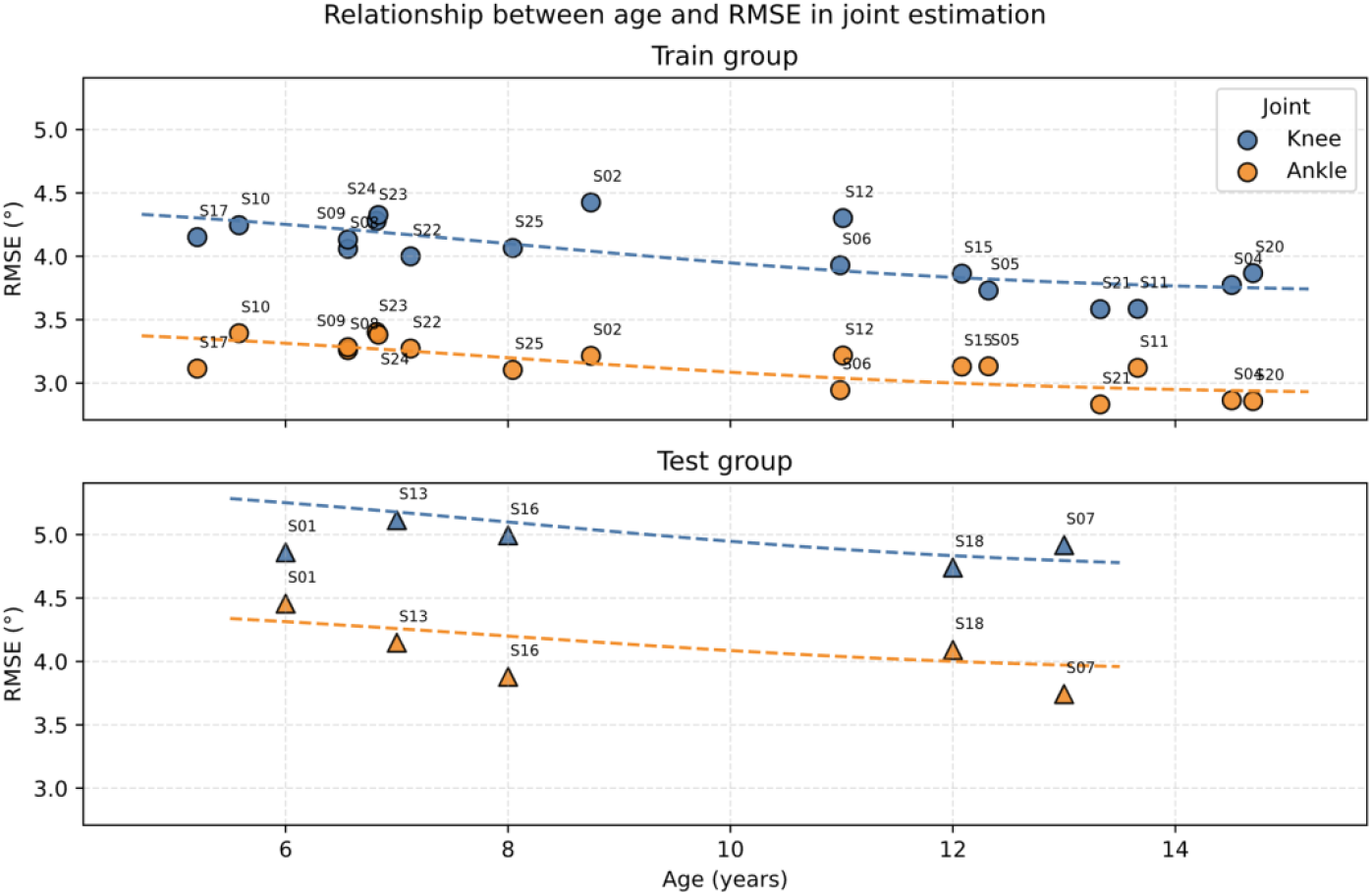
Relationship between age and prediction error in ankle and knee joint estimation. Each point represents one participant.

## 4. Discussion

Synchronization and segmentation between sEMG and IMUs were achieved with high temporal accuracy, yielding mean errors below 0.05 s (≈ 5% of the gait cycle). This precision surpasses that reported in previous studies using neural networks to fuse IMU and sEMG data, which achieved an average error of 1.6% (Caulcrick et al., 2018), (Haufe et al., 2023). Unlike such data-driven approaches, the present method relies on transparent and fully explainable synchronization techniques that can be easily reproduced and adapted to different setups. Therefore, the accuracy obtained in this study is consistent with (and in some cases superior) to state-of-the-art results, confirming the robustness and practical applicability of the proposed synchronization strategy for pediatric gait analysis.

Building on this robust synchronization, analysis of model performance revealed clear differences depending on the joint evaluated, with consistently better outcomes for the ankle compared to the knee. Ankle angle prediction yielded an RMSE of 3.2°, well below the commonly accepted clinical threshold of 5° for gait analysis (Berner et al., 2020), (Haji Hassani et al., 2022). These low errors indicate that the model can reliably reproduce joint trajectories, capturing functional phases and compensatory patterns with high fidelity. High R^2^ values further confirm that predicted signals closely follow the temporal evolution of true joint angles. These results align with previous studies reporting typical ankle errors between 3° and 5° under controlled conditions (Amrani El Yaakoubi et al., 2023; Guez et al., 2025; Wei et al., 2021). Achieving such accuracy using only two sEMG channels, represents a substantial improvement in terms of simplicity, portability, and clinical applicability. The superior ankle performance can be explained by the direct involvement of the TA and MG in ankle control; their activation patterns are closely synchronized with key gait events, allowing the model to learn clear, physiologically grounded relationships between muscle activity and joint motion (Lee et al., 2019). In contrast, knee motion is largely governed by muscles not included in the model (quadriceps and hamstrings) (Frigo et al., 2010), requiring the network to infer behavior indirectly, particularly during mechanically complex transitions, which reduces prediction precision.

The inclusion of TO event as an additional input further enhanced model performance for both joints, providing explicit information about the gait phase. For the ankle, RMSE decreased from 4.9° to 3.2°, and R^2^ increased from 0.8 to 0.9. By incorporating gait phase data, the model can disambiguate muscle activations that may appear similar but serve different biomechanical functions depending on timing. This is particularly relevant for muscles such as the TA, whose activation patterns vary across the gait cycle, contributing to distinct functional roles: before TO to initiate ankle dorsiflexion, during terminal swing to ensure foot clearance, and occasionally in mid-stance to assist in foot inversion control (Di Nardo et al., 2014) (Di Nardo et al., 2013).

Further insights were gained through subgroup analysis by limb dominance, which evaluated the model’s robustness under different neuromuscular control conditions. Prediction errors remained low and well balanced between sides, with slightly better performance on the dominant limb, particularly at the ankle. Although splitting the dataset by dominance effectively halved the number of samples per group, slightly limiting generalization, the results indicate that the model can handle subtle differences in coordination and muscle recruitment that exist between dominant and non-dominant limbs (Pinto et al., 2018) (Lecce et al., 2025). The narrow spread of RMSE values and absence of significant outliers further support the consistency and adaptability of the proposed approach across both limbs.

SPM provided a detailed view of the temporal distribution of prediction errors throughout the gait cycle, revealing that most significant deviations occurred during transitional phases, such as initial contact (0–10%) and pre-swing to early swing (≈60–70%), where rapid kinematic changes and shifts in muscle coordination increase prediction uncertainty. The inclusion of the TO event markedly reduced these deviations, particularly at the ankle, confirming the benefit of incorporating gait phase information. Similar trends were observed at the knee, where TO input minimized discrepancies during terminal swing. Comparisons between dominant and non-dominant limbs revealed similar temporal error distributions, with the dominant limb showing slightly fewer and narrower clusters of significance, reflecting more consistent neuromuscular coordination.

Despite these encouraging results, several limitations should be acknowledged, which also guide future work. First, the study included only typically developing children, and the methodology has not yet been tested in adults or in children with motor impairments, limiting generalizability. Second, although sensor placement followed standardized protocols, small variations in electrode and IMU positioning could have influenced sEMG and kinematic measurements. Third, the model was trained on a specific dataset, and its performance may differ when applied to other cohorts, acquisition systems, or age ranges. These factors should be considered when interpreting the results and assessing their potential clinical applicability.

## 5. Conclusions

This study demonstrates that a one-dimensional U-Net can reliably estimate ankle and knee joint angles from TA and MG sEMG signals in typical developing children, achieving clinically acceptable accuracy (RMSE ≤ 3.6° for the ankle and ≤ 4.1° for the knee). Including the TO event as an additional input significantly improved predictions by providing phase information, particularly during transitional gait phases. Prediction accuracy increased with age, reflecting the maturation of gait patterns in older children. Overall, these results support the use of sEMG-based models for objective, non-invasive gait assessment in typically developing children. Further work is needed to generalize these results to populations with neuromuscular impairments.

## Acknoledgments

The authors would like to express their sincere gratitude to all the individuals who participated in this study.

## Autorship Contributions

C.F.G. B.C., C.G. and M.M.Z. contributed to the conception and design of the study. C.F.G., B.C. and D.I. collected the data and performed data analysis. C.F.G. C.G. and M.M.Z. abstracted and interpreted the results. C.F.G. and H.S. wrote the first draft and approved the final version of the manuscript. B.C., C.G., H.S., D.I., F.C and M.M.Z. revised the manuscript and approved the final version. B.C., M.M.Z. and F.C. were responsible for funding acquisition.

